# Acute inflammation, mediated by lung neutrophils, confers enhanced protection against *Mycobacterium tuberculosis* infection in mice

**DOI:** 10.1101/2021.01.12.426433

**Authors:** Tucker J. Piergallini, Julia M. Scordo, Paula A. Pino, Jordi B. Torrelles, Joanne Turner

## Abstract

Inflammation plays a crucial role in the control of *Mycobacterium tuberculosis* (*M*.*tb*) infection. In this study, we demonstrate that an inflammatory pulmonary environment at the time of infection mediated by liposaccharide (LPS) treatment in mice confers enhanced protection against *M*.*tb* for up to 6 months post infection. This transient protective inflammatory environment was associated with a neutrophil and monocyte/macrophage influx as well as increased inflammatory cytokines. *In vitro* infection of neutrophils from LPS treated mice demonstrated that LPS neutrophils exhibited increased recognition of *M*.*tb*, and had a greater innate capacity for killing *M*.*tb*. Finally, partial depletion of neutrophils in LPS treated mice showed an increase in *M*.*tb* burden, suggesting neutrophils conferred the enhanced protection observed in LPS treated mice. These results indicate a positive role of an inflammatory environment during initial *M*.*tb* infection, and suggests that acute inflammation at the time of *M*.*tb* infection can positively alter disease outcome.

## Introduction

Tuberculosis (TB) disease, caused by the bacterium *Mycobacterium tuberculosis* (*M*.*tb*) is a global health burden. In 2019, 1.4 million people died from TB, making TB the leading cause of death due to an infectious disease worldwide in that year (1). There is variability in *M*.*tb* infection outcome among individuals, with some maintaining infection in a latent state (*M*.*tb* latency) and others developing active TB disease (1). Comorbidities such as HIV co-infection, diabetes, malnutrition, and aging along with environmental exposure to pulmonary insults (wood burning fire, smoking, etc.) are among some of the factors that can increase one’s susceptibility to develop active TB (1, 2). Systemic inflammation has been associated with each of these (3-8).

Inflammation is typically defined as higher levels of inflammatory cytokines and/or a chemokine-mediated influx of immune cells to tissue sites (9). Infection-induced inflammation can have a negative impact on active TB progression (2, 10-12), defining a delicate balance, with both too much and too little inflammation causing worsening prognosis and outlook via host-induced damage or by leading to inadequate control of *M*.*tb* growth, respectively (13-18). Several studies have suggested that the first events immediately after initial *M*.*tb* infection can alter TB disease outcome (19-22), establishing our hypothesis that the inflammatory state of the lung at the moment of first encounter with *M*.*tb* can dictate long term infection and disease outcomes. Comorbidities of diabetes, malnutrition, and aging, along with environmental exposures occur alongside an increased inflammatory state (3-8), making it feasible that transient (acute) *vs*. constant (chronic) increased basal inflammation may define the course of *M*.*tb* infection in humans.

To determine how acute inflammation can influence *M*.*tb* infection outcome, we utilized short-term low dose lipopolysaccharide (LPS) treatment in mice to generate an increased acute systemic and pulmonary inflammatory response at the time of *M*.*tb* infection. LPS treatment caused an increase of inflammatory cytokines and myeloid cells, primarily neutrophils and monocyte/macrophages, in the mouse lungs. Following *M*.*tb* infection, LPS-treated mice had a significant reduction of *M*.*tb* burden evident as early as 7 days post infection, an effect that persisted for at least 6 months. *In vitro* analyses implicated neutrophils as the mediators of early *M*.*tb* control, confirmed through *in vivo* depletion of neutrophils in LPS treated mice prior to *M*.*tb* infection. Our findings confirm that a transient, acute, increased inflammatory environment at the time of *M*.*tb* entry into the lung can impact the course of infection by reducing *M*.*tb* burden in the lung, thus, adding further information to discern why *M*.*tb* exposed people have different infection and disease outcomes.

## Methods

### Mice

Specific pathogen-free male or female BALB/c and male C57BL/6 mice, 11-12 week old, were purchased from Charles River Laboratories (Wilmington, MA) or The Jackson Laboratory (Bar Harbor, ME). Male and female BALB/c mice were used interchangeably as indicated in the figure legends. Mice were housed in animal biosafety level (ABSL) 2 or ABSL3 facilities, in individually ventilated cages, and given sterilized water and standard chow, *ad libitum*. Experimental and control mice were housed in separate cages, and were acclimatized for at least 1 week before use in experiments. Mice were euthanized by CO_2_ asphyxiation. All procedures were approved by the Texas Biomedical Research Institute Institutional Laboratory Animal Care and Use Committee (IACUC), protocol # 1608 MU.

### LPS Mouse Model

Lipopolysaccharides (LPS) (L3129-100MG, Sigma), were injected intraperitoneally (IP) with 20 µg/mouse (BALB/c) or 50 µg/mouse (C57BL/6) in 100 µl normal saline (vehicle). Saline injections served as control. The optimum LPS dose for each individual mouse strain to result in inflammation without persistent morbidity was determined, and mice were injected every 24 hours (hr) ± 2 hr for 4 total injections. Experiments termed day 0 took place 2 hr after the fourth injection. Alternatively, mice were infected with *M*.*tb* 2-4 hr after the fourth injection, as described below. *M*.*tb* infected mice were daily injected with LPS or saline for 2 more days, for a total of 6 injections, and rested until the indicated timepoint.

### *M.tb* stocks

*M.tb* Erdman (ATCC 35801) was obtained from the American Type Culture Collection (Manassas, VA). GFP-expressing *M.tb* Erdman was kindly provided by Horwitz and colleagues (23). Stocks were grown and delivered via aerosol as previously described (24). For *in vitro* infections, a frozen *M.tb* stock was plated onto 7H11 agar (Difco and BBL) supplemented with oleic acid-albumin-dextrose-catalase (OADC) enrichment, and incubated for 11-13 days at 37°C. A *M.tb* single bacterial suspension was generated and diluted to working concentration as described (25).

### *M.tb* aerosol infection and CFU calculation

Mice were exposed to a low-dose aerosol of *M.tb* Erdman using an inhalation exposure system (Glas-col) calibrated to deliver 10-30 colony-forming-units (CFUs) to the lungs of each individual mouse (26). Calculation of *M.tb* (CFU) burden at the indicated timepoints was performed by plating serial dilutions of whole or partial (superior, middle, inferior, and post-caudal lobes) lung homogenates onto OADC supplemented 7H11 agar containing Mycobacteria Selectatab (Mast Group, UK). Plates were incubated at 37°C and CFUs counted after 14-21 days, and transformed to a log_10_ scale. CFU counts obtained from partial lung homogenates were normalized to the mass of each partial lung.

### Protein ELISA and Luminex Analysis

Organ homogenates were thawed and the resulting supernatants analyzed for cytokines and proteins by ELISA (Biolegend, BD, and R&D) and Luminex (R&D) according to manufacturer’s instructions. Protein levels were normalized to lung mass.

### Lung cell isolation

As described previously, mice were euthanized and lungs perfused with 10 ml PBS containing 50 U/ml heparin (Sigma) and placed into 2 ml complete DMEM (c-DMEM); DMEM (10-017-CV, Corning), 500 ml supplemented with filter-sterilized 5 ml HEPES buffer (1 M; Sigma), 10 ml MEM nonessential amino acid solution (100x; Sigma), 5 ml Penicillin-Streptomycin (pen./strep.) (100x; Sigma), 660 µl 2-mercaptoethanol (50 mM; Sigma), and 45 ml heat-inactivated fetal bovine serum (FBS) (Atlas Biologicals). A single-cell suspension was obtained using enzymatic digestion (15). Residual erythrocytes were lysed using Gey’s solution (8 mM NH_4_Cl, 5 mM KHCO_3_ in water), passed through a 40 µm strainer, and suspended in c-DMEM. Total number of viable cells (via.cells) were determined with acridine orange and propidium iodide (AO/PI) staining and counted on a Cellometer K2 Cell Counter (Nexcelom Bioscience).

### Flow cytometry

Lung single-cell suspensions were washed once with 1x Dulbecco’s phosphate buffered saline without calcium or magnesium (Gibco) (PBS) and stained with Zombie Aqua Fixable Viability Kit (1:100; Biolegend) for 15 min at room temperature (RT) in the dark. After incubation, cells were washed once with 1x PBS and incubated in TruStain FcX-Fc Block (Biolegend) for 15 min at 4°C in the dark. Fc Block was diluted in deficient RPMI (dRPMI; RPMI-1640 supplemented with HEPES and 1g/L sodium azide [ThermoFisher Scientific]) with 10% heat-inactivated FBS (dRPMI+FBS). Fc block was then removed and cells incubated in antibody cocktails (CD45.2, PerCP/Cyanine5.5, clone 104; Ly6G, Alexa Fluor 488, clone 1A8; CD11b, APC/Cyanine7, clone M1/70; CD11c, APC, clone N418; Siglec-F, Brilliant Violet 421, clone S17007L; CD80, PE, clone 16-10A1; Biolegend) diluted in dRPMI+FBS for 20 min at 4°C in the dark. After staining, cells were washed once in dRMPI+FBS. For experiments completed at day 0 (uninfected mice), cells were fixed in 2% paraformaldehyde (PFA) for 15 min at RT in the dark, and washed once more in dRPMI+FBS. For experiments completed after infection, cells were fixed in 4% PFA for 30 min at RT in the dark for ABSL3 removal, and washed twice in dRPMI+FBS. All stained and fixed cells were suspended in dRPMI+FBS and stored at 4°C in the dark until data acquisition on a Beckman Coulter CyAn flow cytometer. Data analysis was performed using FlowJo v10 (BD). Total number of specific cell populations per lung calculated using the percentage of each population in the parent gate for absolute quantification multiplied by the total number of viable cells isolated from each sample.

### Collection of adherent lung cells

Lung single-cell suspensions were incubated in tissue-culture plates for 1 hr at 37°C, 5% CO_2_ (15). Plates were washed with c-DMEM to remove non-adherent cells. For RNA isolation, Trizol (ThermoFisher Scientific), was added to the plates and vigorously pipetted. The Trizol solution (containing adherent cell RNA) was frozen at −80°C until RNA extraction. For flow cytometric analysis, adherent cells were incubated in Trypsin-EDTA (Sigma) for 15 min at 37°C, 5% CO_2_, and c-DMEM was added to stop the reaction. Adherent cells were pooled, suspended in c-DMEM and via.cells determined with AO/PI staining as described above. Flow cytometric analysis was performed as described above.

### Real-time PCR

Frozen Trizol samples were thawed, RNA extracted with chloroform, precipitated using isopropanol and 75% ethanol, and reconstituted in DNase/RNase-free water as previously described (15). cDNA was synthesized with random hexamers using an Omniscript RT Kit (Qiagen). cDNA was quantified using TaqMan gene expression probes (ThermoFisher Scientific) and data collected using an Applied Biosystems 7500 real-time PCR instrument. The ΔΔCT method was used to quantify relative numerical units (RNU), normalized to endogenous 18S RNA, relative to saline.

### Isolation of purified lung monocyte/macrophage and neutrophil populations

Lung single cell suspensions were prepared from non-infected mice as described above, with the exception that residual erythrocytes were not lysed with Gey’s solution. For isolation of CD11b^+^ monocyte/macrophages, lung suspensions from 2 saline injected mice were pooled, whereas LPS lung suspensions were individually processed. As CD11b is also highly expressed on neutrophils and natural killer (NK) cells (27, 28), we developed a method which enriched lung single-cell suspensions for monocyte/macrophages by depleting the suspensions of Ly6G^+^ cells (neutrophils) and CD49b^+^ cells (NK cells) via positive selection, followed with positive CD11b magnetic isolation to obtain the leftover monocyte/macrophages. Briefly, suspensions were washed once with selection buffer (1x PBS containing 2% FBS, 1 mM EDTA, and 1x pen./strep.), suspended at a concentration of 1×10^8^ via.cells/ml in 5 ml polypropylene tubes, and first depleted of neutrophils and NK cells using positive magnetic selection, with all incubations carried out in selection buffer using the MojoSort Mouse Ly6G Selection Kit (Biolegend), the EasySep Biotin Positive Selection Kit II (Stemcell Technologies), and biotinylated anti-mouse CD49b antibody (25 µg/ 1×10^8^ via.cells, clone DX5, Biolegend) per each kits’ instructions. Following Fc block incubation (Stem Cell), cells were incubated in Ly6G-selection antibody and CD49b antibody together, followed by the Stem Cell selection cocktail. Finally, cells were incubated with Ly6G and Stem Cell selection magnetic beads together, and selected out using magnets. From the resulting cell suspensions (depleted of neutrophils and NK cells), CD11b^+^ monocyte/macrophages were isolated using positive magnetic selection using the EasySep Mouse CD11b Positive Selection Kit II (Stem cell technologies), according to manufacturer’s instructions.

For isolation of lung neutrophils, single cell suspensions were washed in selection media and suspended at a concentration of 1×10^8^ via.cells/ml. Cells were incubated with normal rat serum (50 µl/ 1×10^8^ via.cells, Stem Cell Technologies and Jackson ImmunoResearch Laboratories) for 5 min. at RT. Neutrophils were isolated using the MojoSort Mouse Ly6G Selection Kit (Biolegend) according to manufacturer’s instructions.

Purified monocyte/macrophage or neutrophil suspensions were washed and suspended in 100-200 µl c-DMEM, and total via.cells determined with AO/PI staining. Cells were washed with 10 ml antibiotic-free c-DMEM (ABFc-DMEM; c-DMEM without pen./strep. added), and suspended at a final concentration of 50,000 via.cells/150 µl in ABFc-DMEM.

### *In vitro M.tb* infection of monocyte/macrophages and neutrophils, and CFU determination

96-well plates and 8-well chamberslides were coated with poly-d-lysine (0.1 mg/ml, Gibco) overnight at RT or 2 hr 37°C, washed three times with 1x PBS, and dried before use. Purified cells were plated at 50,000 via.cells per well in 150 µl ABFc-DMEM in a pre-coated 96 well plate or 8 −well chamberslide. Cells adhered to the plate or slide by centrifuging at 300 x*g*, 5 min, 4°C. A 50 µl single-cell suspension of GFP-*M.tb* Erdman in ABFc-DMEM, calibrated to deliver *M.tb* at a MOI of 5:1, was then added to the wells.

For monocyte/macrophages, samples were infected in duplicate. The infection proceeded for 2 hr at 37°C, 5% CO_2_ (30 minute [min] shaking followed by 1.5 hr of static incubation). After the 2 hr infection, cells were washed three times with ABFc-DMEM, and incubated at static conditions in ABFc-DMEM until the indicated timepoint. For CFU enumeration, cells were centrifuged for 300 x*g*, 5 min, 4°C, and washed three times with ABFc-DMEM, liquid was removed after the final wash. Cold distilled water (50 µl) containing 500 µg/ml DNase I (Sigma) was added and incubated at RT with periodic agitation. 100 µl OADC supplemented 7H9 (Difco) media and 60 µL of 0.25% sodium dodecyl sulfate (Fisher) in 1x PBS were added and cells incubated for 10 min at RT with periodic agitation. 75 µl 20% bovine serum albumin (BSA) (Alfa Aesar) in 1x PBS was added, and wells mixed by pipetting vigorously several times. Resulting solutions were serially-diluted, plated, and incubated as described above. Average CFU expressed as CFU/ml.

For neutrophils, samples were infected in triplicate. The infection proceeded for 30 min at 37°C, 5% CO_2_ with constant shaking. After the 30 min infection, cells were washed three times with ABFc-DMEM, and incubated at static conditions in ABFc-DMEM until the indicated timepoint. For CFU enumeration, cells were centrifuged at 300 x*g*, 5 min, 4°C, and washed three times with ABFc-DMEM, liquid was removed after the final wash. 100 µl 0.1% TritonX-100 (Fisher) in 1x PBS was added, and cells incubated at RT for 15 min with periodic agitation. 100 µl OADC supplemented 7H9 media was added, and wells mixed by pipetting vigorously several times. Resulting solutions were serially-diluted, plated, and incubated as described above. Average CFUs expressed as CFU/ml. For microscopy studies, cells in 8-well chamberslides were washed three times in ABFc-DMEM following the 30 min infection, and then fixed in 4% PFA for 15 min, prior to ABSL3 removal and further processing.

### Immunocytochemistry (ICC) of *in vitro* infections

Fixed cells on chamberslides were washed three times with 1x PBS and stored at 4°C in the dark until staining. Cells were permeabilized by incubating in 0.5% TritonX-100 for 1 min at RT, and washed three times with 1x PBS, 1 min per wash. Blocking buffer (1x PBS with 10% normal donkey serum [Jackson ImmunoResearch Laboratories], 10 mg/ml BSA, and 0.1% Triton X-100) was added and cells incubated for 30 min at 37°C in a humid chamber. Primary antibodies: goat anti-human/mouse myeloperoxidase (1:100, AF3667, R&D) and rabbit anti-mouse histone H3 (citrulline R2 + R8 + R17) (1:200, ab5103, Abcam), diluted in blocking buffer were added and incubated for 1 hr, at 37°C in a humid chamber. Chamberslides were washed three times with 1x PBS, 1 min per wash. Secondary antibodies: donkey anti-goat IgG Alexa Fluor 647 (1:10,000, A21447, ThermoFisher Scientific) and donkey anti-rabbit IgG Alexa Fluor 568 (1:1,000, A10042, ThermoFisher Scientific), diluted in blocking buffer were added and cells incubated for 1 hr at 37°C in a humid chamber. Cells were washed as before with three 1 min washes. Chamberslides were then incubated in 4’,6-Diamidino-2-phenylindole dihydrochloride (DAPI) (1:5,000, ThermoFisher Scientific) in 1x PBS for 5 min, and washed three times in 1x PBS, for 5 min with constant shaking. Chamber sides were mounted with coverslips using Prolong Diamond Antifade Mountant (ThermoFisher Scientific), and dried for at least 24 hr prior to imaging. Chamberslides were analyzed using a Zeiss LSM 800 confocal microscope. 19-22 GFP-*M.tb* (488 nm) events were counted per well. Histone H3 (citrulline R2 + R8 + R17) and DAPI-smear colocalizations were used to mark neutrophil extracellular traps (NETs), and MPO and circular DAPI were used to mark intact neutrophils. In a single-blinded manner, the location of GFP-*M.tb* events were visually assayed as *M.tb* co-localized with an intact neutrophil, or *M.tb* co-localized with a neutrophil NET. Free *M.tb* (outside of cells or NETS) was also determined and no differences were observed between groups. Data are presented as percent fold change of GFP-*M.tb* colocalized with an intact neutrophil or neutrophil NET, relative to the average saline value of each experiment.

### Neutrophil depletion

Anti-mouse Ly6G (300 µg clone IA8, BP0075-1, BioXCell) or its isotype control rat IgG2a (clone 2A3, BP0089, BioXCell) in 1x PBS were administered via the IP route to LPS/saline mice. Antibody injections began the same day as LPS/saline injections, and continued every 48 hr. CFUs were assessed at 7 days of infection. Single cell suspensions were isolated at day 0 (uninfected) and analyzed by flow cytometry to assess depletion efficiency. Intracellular Ly6G was used in place of surface Ly6G to account for potential surface antigen masking by the depletion antibody (29). After 2% PFA fixation, intracellular Ly6G was stained in the Intracellular Staining Permeabilization Wash Buffer (Biolegend) using the manufacture’s protocol. Ly6G antibodies (Ly6G, PE, clone 1A8, BD Pharmingen) used for intracellular staining were diluted half of what was typically used for surface staining.

### Statistical analysis

Data analyses, graphing, and statistical analyses were performed using GraphPad Prism 8 and 9 software. Unpaired, two-tailed Student’s *t*-test was used for two group comparisons. Statistical significance is reported as \**p*<0.05; \*\**p*<0.01; \*\*\**p*<0.001, or ****p<0.0001. The Grubbs’ test was used to identify outlying data points. Data are presented as individual data points, and mean ± SEM.

## Results

### LPS causes pulmonary and systemically increased inflammation in mice

To evaluate the impact of acute inflammation on *M.tb* infection, we injected BALB/c mice with LPS or saline via the IP route every 24 hr, for 4 total injections (Fig. 1A). Two hr after the fourth injection, termed day 0, we analyzed the local inflammatory response in the lungs. TNF, IL-1β, IL-6, IL-12p70, and IL-10 were significantly increased in the lungs of LPS-injected mice (LPS mice) (Fig. 1B). We also saw a concomitant increase of TNF, IL-1β, IL-6, and IL-10 in the spleens of LPS mice, while IL-12p70 showed no differences (Supp. Fig. S1A). C-reactive protein (CRP) levels showed no difference in the lung (Supp. Fig. S1B), but was increased in spleen and liver (Supp. Fig. S1B).

**Figure 1.**
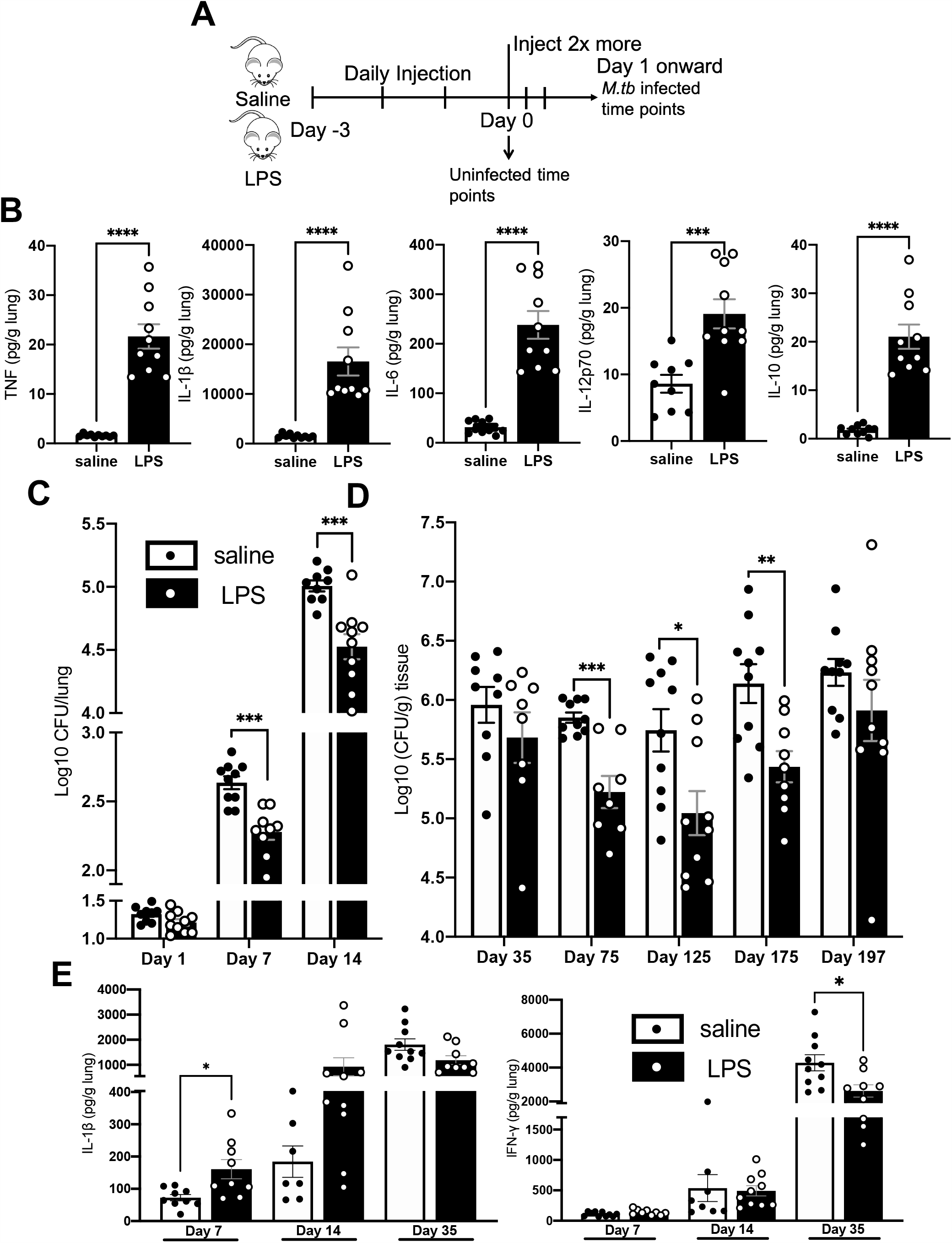
Cytokines and CFUs in LPS mice. **A**. Schematic of injection scheme **B**. Male BALB/c mice were injected with LPS as described. On Day 0, lungs were isolated, and protein content determined via Luminex, normalized to organ mass. Protein levels of TNF, IL-1β, IL-6, IL-12p70, and IL-10 are shown. **C**.**D**. LPS or saline BALB/c mice were aerosol-infected with *M.tb* as described. CFU burden at the indicated timepoint of whole (C) or partial (D) lungs from male (C) and female (D) mice are shown. Partial lung CFU were normalized to lung mass. **E**. At the indicated timepoint, protein levels via ELISA of IL-1β and IFN-γ in male BALB/c mice are shown. Data are representative of 2 independent experiments of 4 or 5 mice in each group. unpaired Student’s t test, *P<0.05, **P<0.01, ***P<0.001, ****P<0.0001.

### Acute inflammation protects mice from M.tb infection

We infected LPS mice with *M.tb* on day 0 (4 days after initiation of LPS/saline treatment), injected with LPS/saline daily for 2 more days, and rested until the indicated timepoints (Fig. 1A). At 1 day post infection (d.p.i.), we found no difference in CFU (Fig. 1C), whereas at 7 and 14 d.p.i. we saw significantly fewer CFUs (∼0.5 log_10_) in LPS mice (Fig. 1C). To determine if the early control of *M.tb* was not mouse strain specific, we repeated these studies in C57BL/6 mice, showing that at 14 d.p.i., C57BL/6 treated LPS mice also had significantly fewer CFU compared to saline-injected mice (saline mice) (Supp. Fig. S1C). These results suggest that the effects of LPS on the early control of *M.tb* was not mouse strain specific, but was related to LPS induced inflammation. We further determined the CFU content at 35, 75, 125, 175, and 197 d.p.i., and observed lower CFU levels (∼0.25, ∼0.6, ∼1.0, ∼1.0, ∼0.4 log_10_ [CFU/g tissue], respectively) in LPS mice at all timepoints (Fig. 1D), although only days 75, 125, and 175 showed statistical significance.

When determining cytokine protein levels at 7, 14, and 35 d.p.i., we observed that IL-1β levels in the lung were elevated at 7 d.p.i., with a trend increase at 14 d.p.i. (Fig. 1E). IFN-γ levels in lung showed no differences at 7 and 14 d.p.i. (Fig. 1E). TNF levels showed no differences at 7 d.p.i., but showed a trend increase at 14 d.p.i. when compared to saline (Supp. Fig. S1D). At 35 d.p.i., LPS mouse lungs showed significantly less IFN-γ, and a trend decrease of IL-1β and TNF, likely a result of their lower level of *M.tb* CFU burden at this time when compared to saline treated *M.tb* infected mice (Fig. 1E and Supp. Fig. S1D).

### LPS mice have more activated myeloid cells in the lungs

To determine the cellular mechanism behind the early *M.tb* control in LPS mice, we analyzed lung cellular profiles at day 0, prior to infection. LPS mice had more viable cells (Fig. 2A), and more myeloid cells (CD45^+^SSC^hi^) per lung (Fig. 2B). Flow gating schemes and representative images of LPS or saline treated mouse profiles are shown in Supp. Fig. S2. Further analysis showed that LPS mice had higher amounts of neutrophils (CD11b^+^Ly6G^hi^) (Fig. 2C), alveolar macrophages (AMs) (Ly6G^lo/neg^CD11c^+^SiglecF^+^) (Fig. 2D), and monocyte/macrophages (mon./mac.), which both singularly expressed CD11b (Ly6G^lo/neg^CD11b^+^CD11c^-^) (Fig. 2E) and dually expressed CD11b and CD11c (Ly6G^lo/neg^CD11b^+^CD11c^+^) (Fig. 2F) in LPS mice. Eosinophil (eos.) (Ly6G^lo/neg^CD11c^+^SiglecF^-^) numbers were relatively unchanged (Fig. 2G). CD80 is upregulated on macrophages after activation (30), and we found higher numbers of CD80^+^ CD11b^+^CD11c^-^ mon./mac., and CD80^+^ CD11b^+^CD11c^+^ mon./mac (Fig. 2 H,I) in LPS mice. No differences were seen in CD80^+^ AMs (Fig. 2J). We also determined CD11b expression on neutrophils and CD11b^+^CD11c^-^ mon./mac and found a significant lower level of CD11b mean fluorescence intensity (MFI) on neutrophils, and a significant higher CD11b MFI on mon./mac. (Fig. 2 K,L) in LPS mice, suggesting higher CD11b surface expression on the mon./mac, and less on neutrophils from LPS mice.

**Figure 2.**
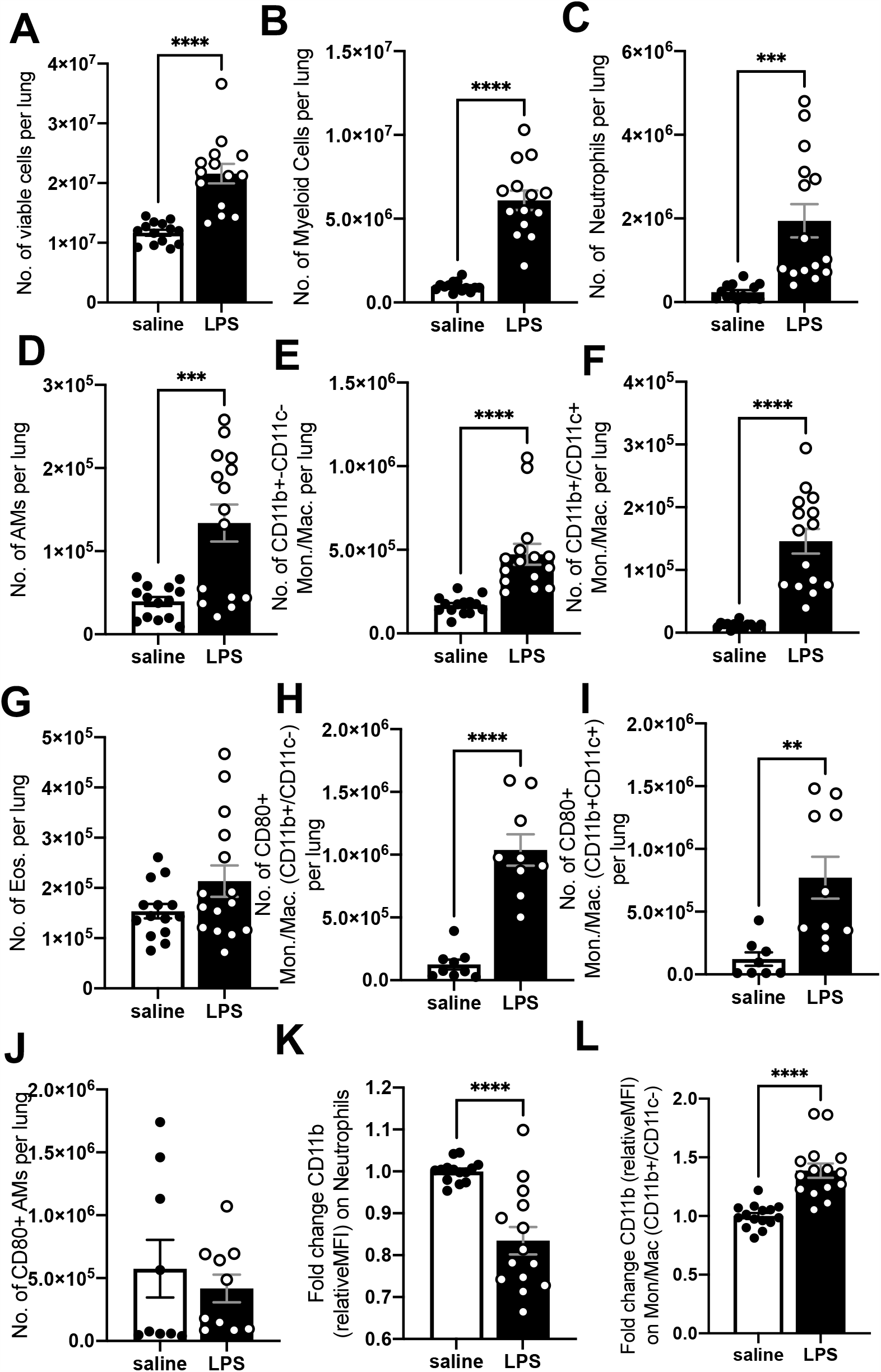
Day 0 lung cell profiles. Single-cell lung suspensions were obtained at day 0 (uninfected) from male BALB/c mice. **A**. Total number of viable cells and **B**. myeloid cells isolated from the lungs of the mice. **B-L**. Flow cytometric analysis of single cell suspensions. Gating done as described in Supp. Fig. S2. **C-G**. Number (No.) of Neutrophils (CD11b^+^Ly6G^+^) (C), Alveolar macrophages (AMs) (Ly6G^lo/neg^CD11c^+^SiglecF^+^) (D), CD11b^+^ monocyte/macrophages (mon./mac.) (Ly6G^lo/neg^CD11b^+^CD11c^-^) (E), CD11c^+^ mon./mac. (Ly6G^lo/neg^CD11b^+^CD11c^+^) (F), and eosinophils (eos.) (Ly6G^lo/neg^CD11c^-^SiglecF^+^) (G) per lung. Absolute number of cells per lung are shown. **H-J**. No. of CD80^+^ CD11b^+^ mon./mac. (H), CD11c^+^ mon./mac. (I), and AMs (J) per lung. **K-L**. Relative fold change CD11b expression (MFI) on neutrophils (K) and CD11b^+^ mon./mac. (L), relative to saline. Data are representative of 2-3 independent experiments with 4-5 mice in each group.; unpaired Student’s t test, **P<0.01, ***P<0.001, ****P<0.0001.

### Elevated numbers of myeloid cells in LPS mice persist at 1 week, but normalize by 2 weeks post *M.tb* infection

We next determined when, and if, the levels of myeloid cells normalized in *M.tb* infected LPS mice relative to *M.tb* infected saline mice. Total viable cells in *M.tb* infected LPS mice were elevated at 7 d.p.i., but no differences were seen between groups at 14 d.p.i. (Fig. 3A). The same trend was observed in total myeloid cells at 7 and 14 d.p.i. (Fig. 3B). Flow cytometric analysis of specific myeloid cell populations showed higher numbers of neutrophils (Fig. 3C), CD11b^+^CD11c^-^ mon./mac. (Fig. 3D), and CD11b^+^CD11c^+^ mon./mac. (Fig. 3E) in *M.tb* infected LPS mice at 7 d.p.i., which normalized to *M.tb* infected saline mice by 14 d.p.i. AMs (Fig. 3F) and eos. (Fig. 3G) showed no significant differences at 7 or 14 d.p.i. Elevated cell numbers in LPS mice compared to saline mice at 7 d.p.i., but not 14 d.p.i., indicated that the inflammatory stimulus from LPS treatment in *M.tb* infected LPS mice was acute. Furthermore, we observed CD11b MFI differences for both neutrophils and CD11b^+^CD11c^-^ mon./mac at 7 d.p.i., but only the former was significant. At 14 d.p.i., CD11b MFI differences were normalized on neutrophils and CD11b^+^CD11c^-^ mon./mac from LPS and saline mice (Fig. 3H,I). These results show that the *M.tb* infected LPS mice were under acute inflammation at the time of infection, and the cellular events responsible for the early protection against *M.tb* likely occurred within the first week post-infection. This was supported by the lower CFUs in LPS mice as early as 7 d.p.i. (Fig. 1C).

**Figure 3.**
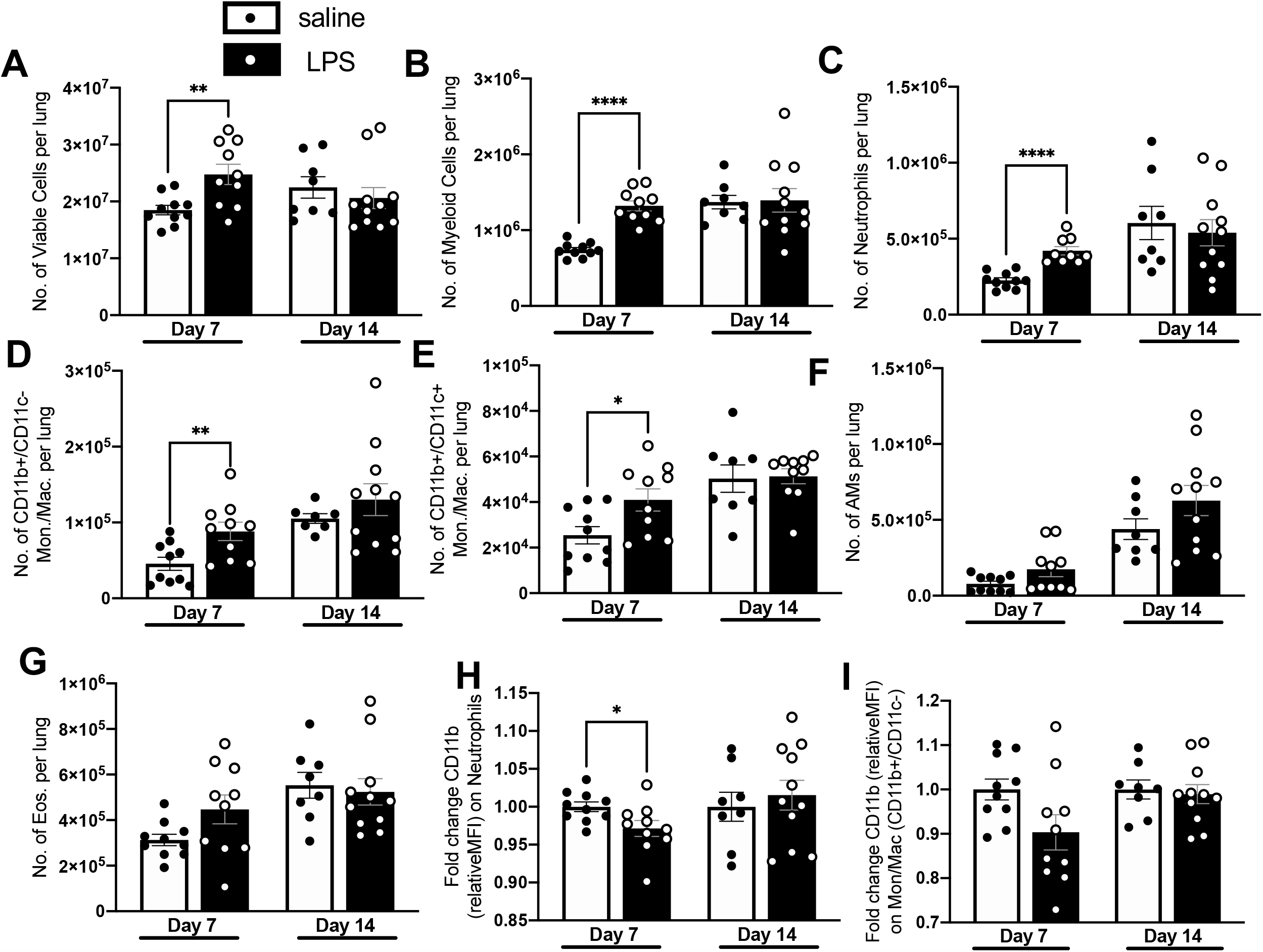
Lung cell profiles at 7 and 14 d.p.i. Lung single cell suspensions prepared at the indicated timepoint post-infection as described from male and female BALB/c mice. **A**. Total number of viable cells and **B**. myeloid cells. **B-L**. Flow cytometric analysis of single cell suspensions. Gating done as described in Supp. Fig. S2. **C-D**. No. of neutrophils (CD11b^+^Ly6G^+^) (C), CD11b^+^ mon./mac. (Ly6G^lo/neg^CD11b^+^CD11c^-^) (D), CD11c^+^ mon./mac. (Ly6G^lo/neg^CD11b^+^CD11c^+^) (E), AMs (Ly6G^lo/neg^CD11c^+^SiglecF^+^) (F), and eos. (Ly6G^lo/neg^CD11c^-^ SiglecF^+^) (G) per lung per lung. **H-I**. Relative fold change CD11b expression (MFI) on neutrophils (H) and CD11b^+^ mon./mac. (I) relative to saline. Data are representative of 2 independent experiments with 4-5 mice in each group. unpaired Student’s t test, *P<0.05, **P<0.01, ****P<0.0001.

### Macrophages from LPS mice do not contribute to *in vitro* control of *M.tb*

To determine the contribution of macrophages in the early *M.tb* control by LPS mice, we measured mRNA expression levels of several inflammatory transcription factors (CIITA, IRF1, and IRGM1) (31-34) from adherent lung cells at day 0 (uninfected), and 7 and 14 d.p.i. Flow cytometric analysis found the majority of adherent cells to be AMs, with some mon./mac. (total adherent cells referred to as pulmonary macrophages) (Supp. Table 1). We observed no differences in CIITA mRNA levels at any timepoint (Fig. 4A) whereas IRF-1, and IRGM-1 showed an increase in *M.tb* infected LPS mice at 7 d.p.i. only (Fig. 4B,C). We concluded that pulmonary macrophages in LPS mice transiently upregulated inflammatory transcription factors after day 0.

**Table 1:** Neutrophils present in mice lungs depleted of neutrophils at day 0.

| Sample | Number of neutrophils ± SD<br>(if applicable) | Fold change vs. saline<br>isotype |
| --- | --- | --- |
| **LPS isotype** | $7.08 \times 10^6$ | 32.32 |
| **LPS depletion** | $3.58 \times 10^6 \pm 4.25 \times 10^5$ | 16.34 |
| **saline isotype** | $2.19 \times 10^5$ | 1.00 |
| **saline depletion** | $1.11 \times 10^5$ | 0.50 |

The average absolute number of neutrophils per lung at day 0 are shown. Saline mice included as reference. Data representative of 1 independent experiment with 1-2 samples in each group.

**Figure 4.**
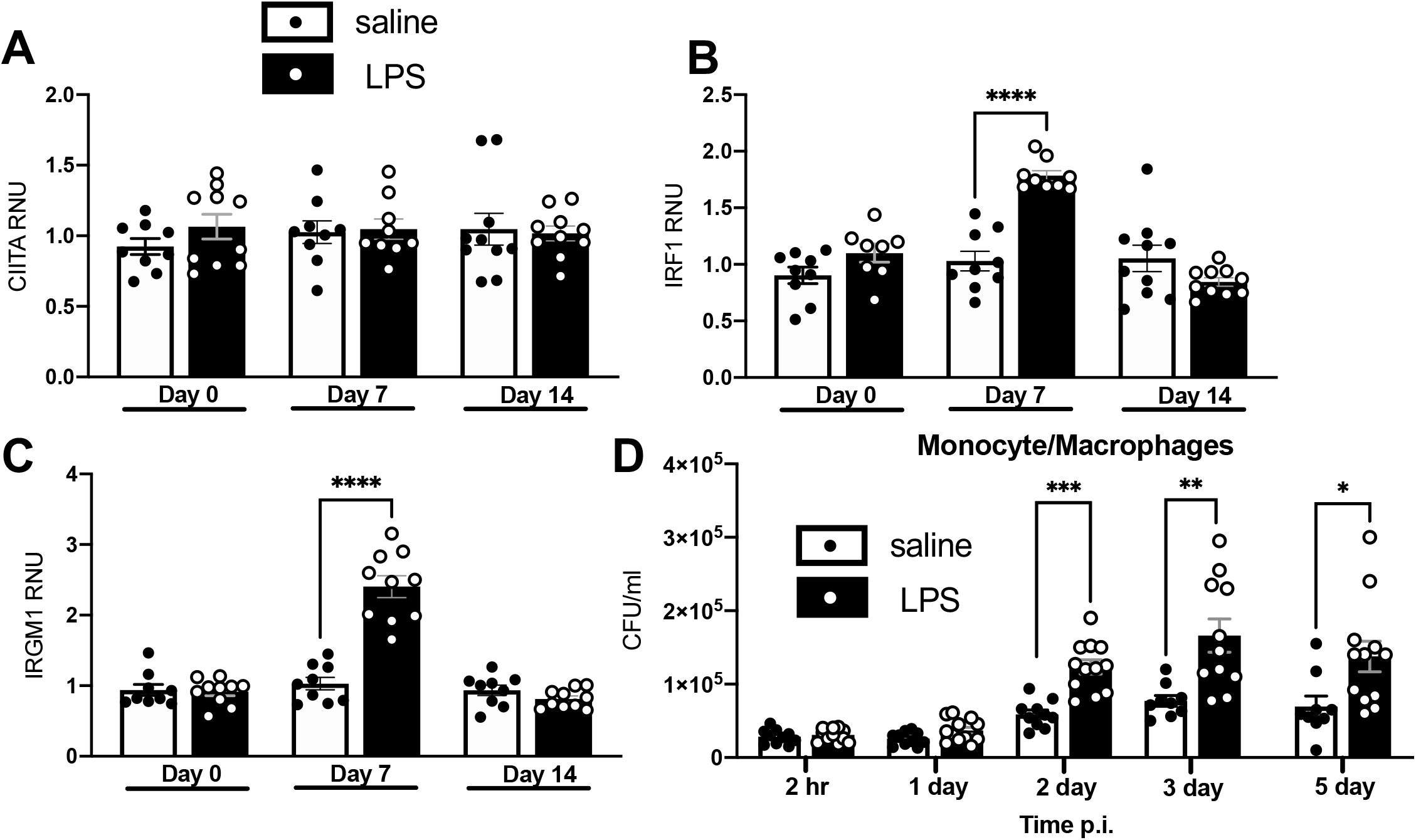
Macrophages from LPS mice are transiently activated, but detrimental to *in vitro M.tb* infection. **A-C**. At the indicated timepoint, RNA was isolated from pulmonary macrophages from female BALB/c mice and quantified using qRT-PCR. CIITA (A), IRF1 (B), and IRMG1 (C) levels are shown. Data is expressed as relative numerical units (RNU) relative to saline mice, and is representative of 2 independent experiments of 4 or 5 mice in each group. **D**. Mon./mac. were isolated from uninfected female BALB/c mice and infected *in vitro* with *M.tb*. CFUs at the indicated timepoint are shown. Data express as CFU/ml and are represented by 3 independent experiments with 2-5 samples in each group. unpaired Student’s t test, *P<0.05, **P<0.01, ***P<0.001, ****P<0.0001.

We next purified mon./mac. at day 0 (uninfected) from LPS or saline mice based on CD11b positive magnetic selection (Supp. Table 1), infected *in vitro* with GFP-expressing *M.tb* Erdman, and determined CFUs at 2 hr, and 1, 2, 3, and 5 d.p.i. We observed no differences between groups at 2 hr and 1 d.p.i., but 2, 3, and 5 d.p.i. showed significantly higher *M.tb* CFUs in mon./mac. isolated from LPS mice (Fig. 4D). This suggested that the mon./mac. infiltrates we saw in LPS mouse lungs at day 0 may not be contributing to the early control we see after *in vivo* infections, and may possibly be detrimental.

### Neutrophils are responsible for enhanced control of *M.tb* infection in LPS mice

LPS treatment also increased neutrophils in the lung (Fig. 2C). To determine the potential involvement of neutrophils in control of *M.tb* infection in LPS mice, we isolated neutrophils via Ly6G positive magnetic selection at day 0 (uninfected) (Supp. Table 1), and infected the purified cells *in vitro* with GFP-*M.tb* Erdman. At 30 min, 1 hr, and 2 hr post infection, we determined CFU (Fig. 5A). While at each timepoint there was no significant difference between CFU in LPS and saline mice, we found paired LPS neutrophils had significantly less *M.tb* CFU at 2 hr relative to 30 min when compared to paired saline neutrophils (Fig. 5B), suggesting a greater magnitude of *M.tb* killing. When we analyzed the infection using single-blinded ICC microscopy (Fig. 5C), we found no differences in *M.tb* association with neutrophil extracellular traps (NETs), but the numbers of *M.tb* co-localizing with intact neutrophils was significantly increased in neutrophils from LPS mice when compared to saline mice (Fig. 5D).

**Figure 5.**
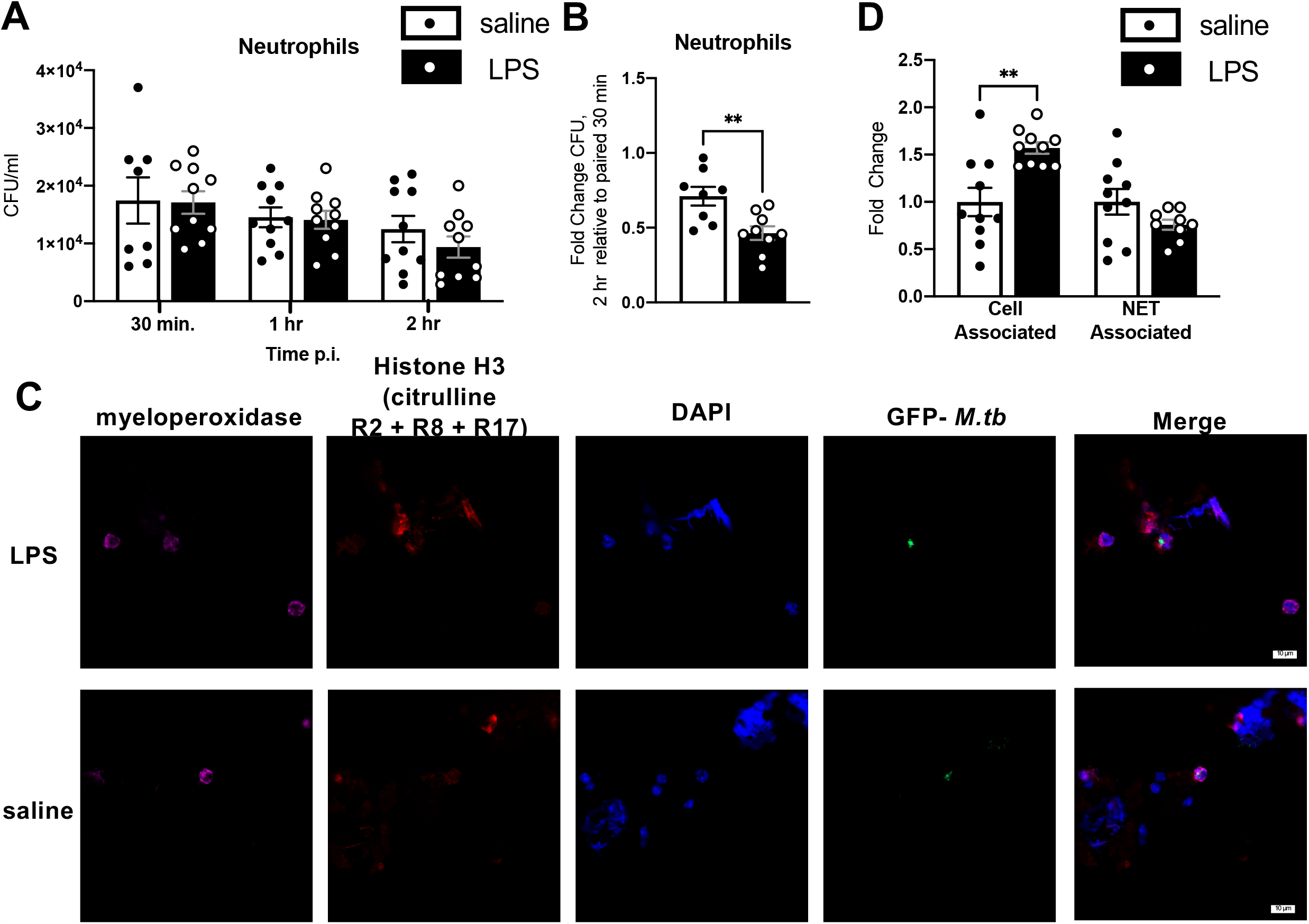
Neutrophils from LPS mice are capable of increased *M.tb* killing. Neutrophils were isolated from uninfected female BALB/c mice as described. **A**. CFUs during *in vitro* infections. **B**. Fold change of CFUs over 2 hr of neutrophil infection, calculated relative to paired 30 min. data. **C**. Immunocytochemistry of neutrophil infections. Pink-myeloperoxidase, red-citrullinated histone H3 (R2+R8+R17), blue-DAPI, green-GFP *M.tb*. Representative images of a GFP-*M.tb* event colocalized with an intact cell (neutrophil) shown. **D**. Percent fold change of the location of GFP-*M.tb* in each well, relative to saline. Slides analyzed in a single-blinded manner. 19-22 GFP-*M.tb* events analyzed per well. Cell-associated, GFP-*M.tb* colocalized with an intact cell; NET associated, GFP-*M.tb* colocalized with a neutrophil NET. Data are representative of 2 independent experiments with 4-5 samples in each group. unpaired Student’s t test, **P<0.01.

To confirm that LPS neutrophils were responsible for conferring increased control of *M.tb*, we depleted neutrophils over the course of *in vivo* infection (Table 1). At 7 d.p.i., *M.tb* infected LPS mice depleted of neutrophils showed higher levels of *M.tb* CFU compared to *M.tb* infected LPS mice injected with the isotype control (Fig. 6), although these results failed to reach statistical significance. Increased killing *in vitro* (Fig. 5), together with these results, point to neutrophils as being the mediator behind increased early control of *M.tb* infection in LPS mice *in vivo*.

**Figure 6.**
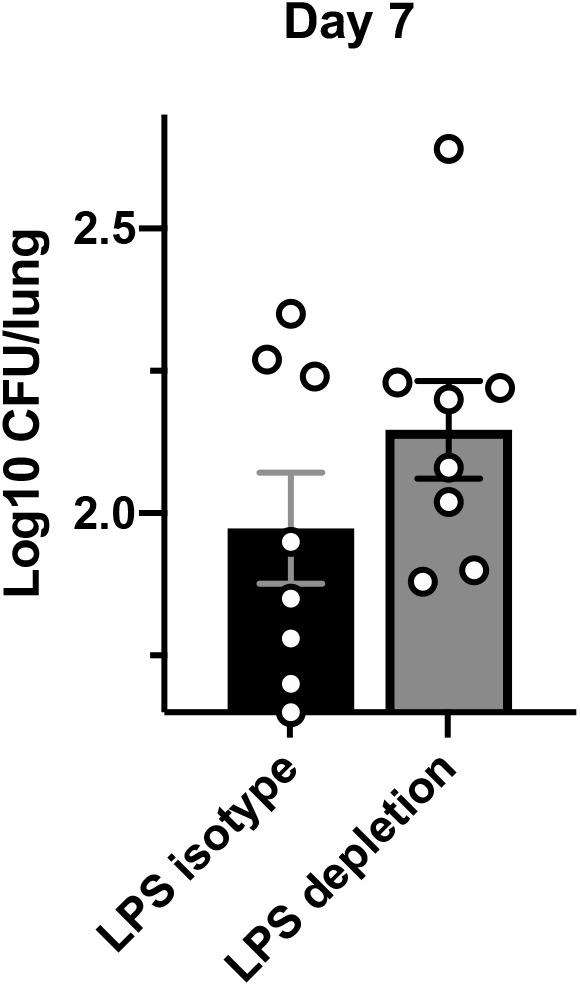
Neutrophils drive control in LPS mice. Neutrophils were depleted from LPS and saline female BALB/c mice using monoclonal antibodies against Ly6G as described. CFUs at 7 d.p.i. in LPS mice injected with depletion or isotype antibodies shown. Data are representative of 3 independent experiments with 2-4 samples in each group. unpaired Student’s t test, results not statistically significant.

## Discussion

Inflammation can affect the control of TB disease (10-12), yet the impact of basally increased inflammation (occurring in the elderly, persons with lung co-infections, those with environmental insult, etc.) at the time of *M.tb* infection has received limited attention (2, 19, 20). To investigate this, we developed a mouse model to test if acute inflammation induced by a short-term low level LPS delivery could alter control of *M.tb* infection. LPS is a potent inflammatory stimulus, and causes an upregulation of many inflammatory cytokines and signals (35, 36). Mice injected with LPS established an acute pulmonary and systemic inflammatory state and an influx of neutrophils and monocyte/macrophages into the lungs prior to infection, which can be expected after an acute inflammatory stimulus (37). After *M.tb* infection, LPS mice had lower *M.tb* burden compared to controls as early as 7 d.p.i., and continuing up to 6 months post infection. *In vitro* and *in vivo* studies demonstrated that neutrophils, and not monocyte/macrophages, were the driver of enhanced control in LPS mice. These findings suggest that acute inflammation, driven by an increase of neutrophils, can confer protection against *M.tb* infection in the mouse model.

Our results suggest that an acute inflammatory state at the time of *M.tb* infection can be protective against *M.tb*. Examples of this in the literature include studies in old mice, which are chronically inflammatory, both in the periphery and the lung (15, 16, 38). This chronic inflammatory status makes old mice display an early control of *M.tb* infection compared to their younger counterparts (21, 39, 40). This early control is considered a consequence of the chronic but moderate inflammatory state at the time of infection in old mice, as work from our group has indicated (14, 41-43), and we showed that innate cells in the lungs of old mice are pre-activated and behave differently in response to *M.tb* (15). In contrast to LPS mice (acute model), old mice (chronic model) cannot sustain control, likely due to their reported reduced adaptive immune function (44, 45), and old mice succumb to the infection earlier than young mice (39, 46). Adaptive immune function was not investigated in LPS mice, but as we observed decreased *M.tb* burden in LPS mice up to 6 months p.i., the longest timepoint we tested, the initial innate response against *M.tb* in LPS mice was enough to maintain protection long term in our model.

In human active TB, neutrophils are the most infected phagocytic cell and can provide a niche for *M.tb* persistence and survival (47). Furthermore, neutrophil influx to the lungs is associated with worse TB disease in patients (48). Neutrophils enter the lung in high numbers after *M.tb* infection, and are typically reported to kill *M.tb* via phagocytosis and subsequent killing, as well as extracellular killing mechanisms (degranulation) (49, 50). Neutrophil NETs can also contribute to control via slowing of *M.tb* growth and *M.tb* death (51), but are also reported to contribute to the worsening of disease (52). The amount of conflicting information on neutrophils in *M.tb* infection suggests the role of neutrophils is context dependent. Indeed, rodent studies show that neutrophils play a beneficial role in early infection, but a negative role at later stages (19, 20, 53, 54). A study of LPS induced lung neutrophilia in rats showed reduced *M.tb* burden if LPS was delivered prior to infection, which was negated following neutrophil depletion (19). LPS delivery 10 days post *M.tb* infection, however, had no effect. Our results from LPS mice corroborate these findings, and suggest that neutrophils can play a beneficial role in early *M.tb* infection, if they are increased in number and primed prior to the arrival of *M.tb* to the lungs. In our own neutrophil depletion studies, however, we recognize that the reduced *M.tb* CFU from our neutrophil depletions of LPS mice did not reach statistical significance, possibly because of incomplete depletion of neutrophils in LPS mice. It is accepted that depletion of neutrophils in mice with the 1A8 clone is difficult for a variety of reasons (29). After depletion, the bone marrow generates neutrophils to maintain homeostasis (55) which, coupled with the LPS stimulus, likely caused abundant neutrophils present in the periphery to be depleted, resulting in incomplete depletion.

The role of neutrophils in mediating control of *M.tb* infection in LPS mice is supported by our *in vivo* flow cytometry data, where neutrophil numbers were increased the most in lungs of LPS mice compared to other cells. Our *in vitro* work also suggests that neutrophils from LPS mice phagocytose and kill *M.tb* more effectively. Furthermore, we observed less CD11b expression on LPS neutrophils, a component of complement receptor 3 (CR3) (56). CR3 is one of the major phagocytic receptors for *M.tb* and can be activated by *M.tb* on neutrophils, although it is not known if neutrophils directly use CR3 to phagocytose *M.tb* (56-58). Because we observed no negative impact in the co-localization of *M.tb* and LPS neutrophils, we can conclude that the lower levels of CD11b expression on LPS neutrophils had a limited effect. Interestingly, a recent experiment showed that lowering levels of CD11b expression on neutrophils resulted in increased protection against *M.tb* in a mouse model, although this was associated with decreased neutrophil accumulation in the lungs (59). Given the complicated and often disparate roles of neutrophils shown in the literature, more studies are needed to address the role of neutrophils in differing states of *M.tb* infection, especially in cases where neutrophil behavior may be altered (19, 20, 53, 54, 59).

The other cell type we interrogated as potentially being responsible for control in LPS mice were monocyte/macrophages, and our results suggested monocyte/macrophages are not major contributors to the increase in *M.tb* control seen LPS mice, at least *in vitro*. As *M.tb* infections progress in a mouse, monocyte/macrophages are known to be essential for infection control, and the transition of *M.tb* to interstitial macrophages has been shown as beneficial (60, 61). However, interstitial macrophages can also provide *M.tb* a niche for survival (62-64). A recent experiment on an alveolar macrophage subset showing monocytic markers in old mice displayed worse control of *M.tb* (13), and in humans and mice infected with *M.tb*, a monocyte influx to the lungs correlates with worse TB disease outcome (65, 66). These substantiate the results from our *in vitro* infections of monocyte/macrophages in LPS mice. However, additional studies are required to assess the interplay of inflammatory monocyte/macrophages and *M.tb* in our LPS mouse model.

Overall, our results demonstrate that basal inflammation at the time of *M.tb* infection can impact infection outcome. In a broader sense, this suggests that inflammation driven by acute insult at the time of *M.tb* infection can lead to improved long term control of infection. This mechanism may have relevance for some co-infections and other disease states that establish a short-term (acute) pulmonary inflammation in the lung that induces neutrophil influx. In these cases, the initial bacterial burden may be lowered during early *M.tb* infection and maintained long-term, giving rise to differential TB infection outcomes.

## Author’s contributions

TJP and JT designed the experiments. TJP and JMS performed the experiments. PAP assisted with mouse procedures. TJP collected and analyzed data, and wrote the paper. JT and JBT provided critical review of the paper. All authors read and approved the final manuscript.

## Acknowledgments

We would like to acknowledge the Texas Biomedical Research Institute Biology Core for their assistance with data collection for flow cytometry and Luminex assays.

## Figure Legends

**Supplementary Figure 1. Cytokines and CFUs in LPS mice A-C**. On Day 0, male BALB/c LPS and saline organs were isolated, and protein content determined via Luminex (A) and ELISA (B). TNF, IL-1β, IL-6, IL-12p70, and IL-10 in spleen (B), and CRP in lung, spleen, and liver (B) and are shown. Data is normalized to organ mass **C**. LPS or saline male C57BL/6 mice were aerosol-infected *M.tb* Erdman. CFU content shown at 14 d.p.i. **D**. LPS or saline male BALB/c mice were aerosol-infected with *M.tb* as described. At the indicated timepoint, protein levels via ELISA of TNF are shown. Data are representative of 2 (A,B,D) or 4 (C) independent experiments of 2-5 mice in each group. unpaired Student’s t test, **P<0.01, ***P<0.001, ****P<0.0001.

**Supplementary Figure 2. Flow Cytometry Gating strategy used. Gating based on fluorescence minus one (FMO) controls. A-H**. Doublets (A) and dead cells (B) gated out. Parent gate used for absolute number quantification (C). **D-H** Representative flow cytometry images from LPS and saline mice. CD45^+^ cells (D), myeloid cells (SSC-L^hi^) (E), neutrophils (CD45^+^SSC-L^hi^CD11b^+^Ly6G^hi^) (F), eosinophils (CD45^+^SSC-L^hi^Ly6G^lo/neg^SiglecF^+^CD11c^-^) and alveolar macrophages (CD45^+^SSC-L^hi^Ly6G^lo/neg^SiglecF^+^CD11c^+^) (G), and monocyte/macrophages (CD45^+^SSC-L^hi^Ly6G^lo/neg^SiglecF^-^ CD11b^+^CD11c^+^) and (CD45^+^SSC-L^hi^Ly6G^lo/neg^SiglecF^-^ CD11b^+^CD11c^-^) (H).

**Supplementary Figure 3. Intracellular flow cytometric analysis of lung neutrophils in mice injected with neutrophil depleting antibody or isotype as described.** Gating strategy used as in supp. Fig. S2. **A**. Representative images of surface Ly6G vs. intracellular Ly6G in LPS mice. (CD45^+^SSC-L^hi^CD11b^+^), gated from total CD11b^+^ myeloid cells. **B**. Representative images of LPS/saline mice injected with depletion antibody/isotype at day 0. Ly6G is stained intracellularly. (CD45^+^SSC-L^hi^CD11b^+^Ly6G^hi-intracellular^), gated from total myeloid cells.

