## Supplemental Figures and Table for "Acute inflammation, mediated by lung neutrophils, confers enhanced protection against *Mycobacterium tuberculosis* infection in mice"

### Supplementary Figure 1. Cytokines and CFUs in LPS mice

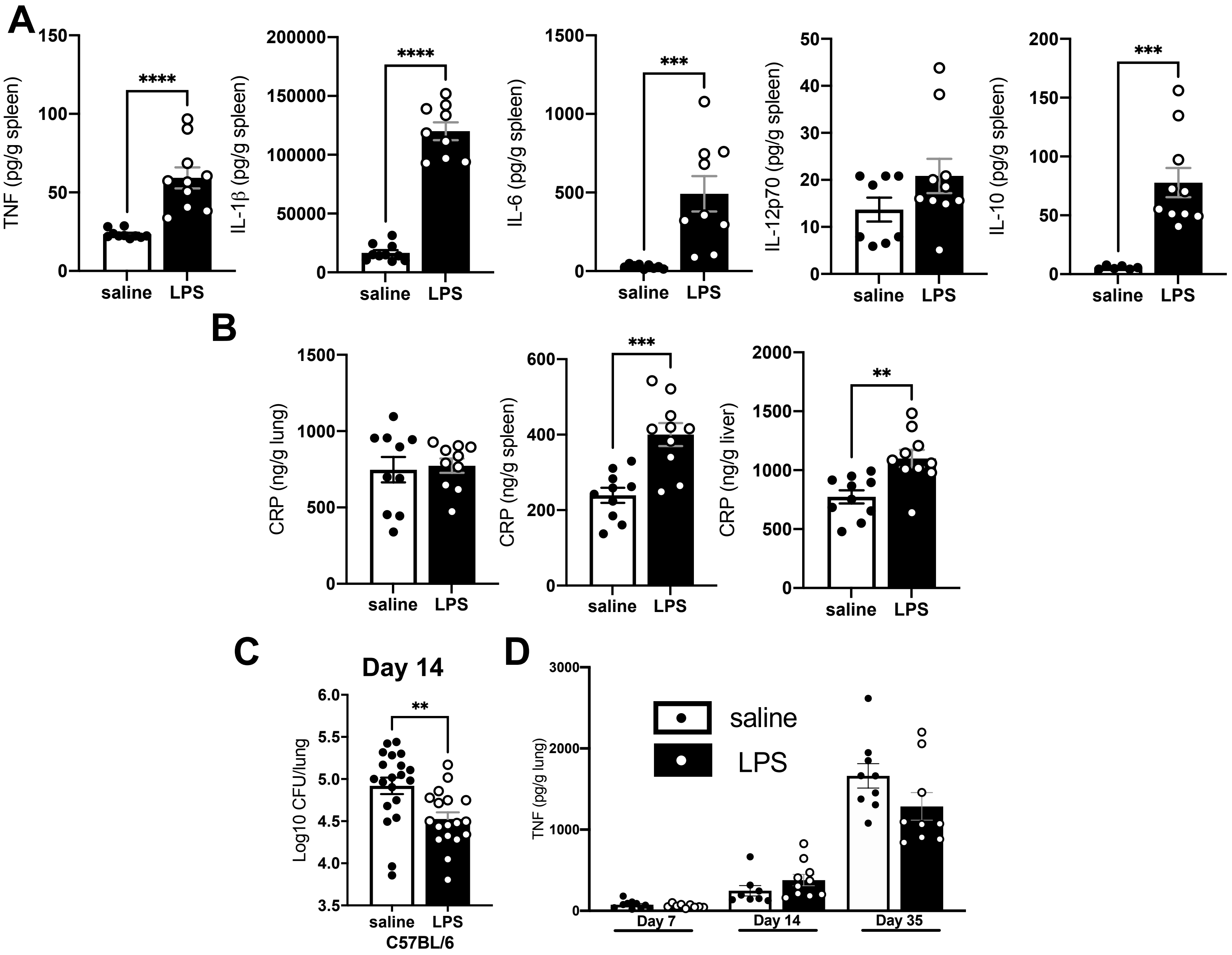

### Supplementary Figure 2. Flow Cytometry Gating strategy used.

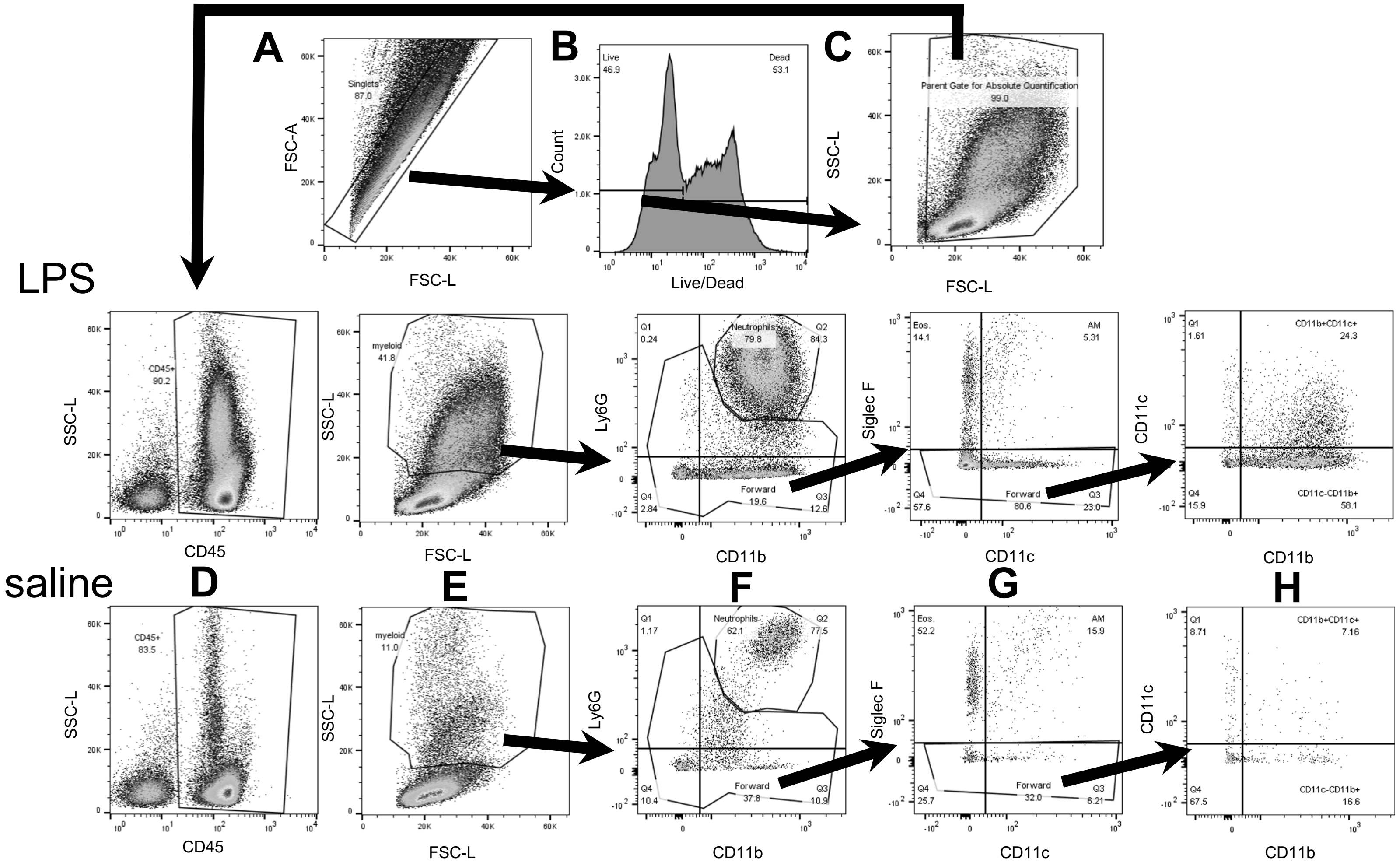

### Supplementary Figure 3. Intracellular flow cytometric analysis of lung neutrophils in mice injected with neutrophil depleting antibody or isotype as described.

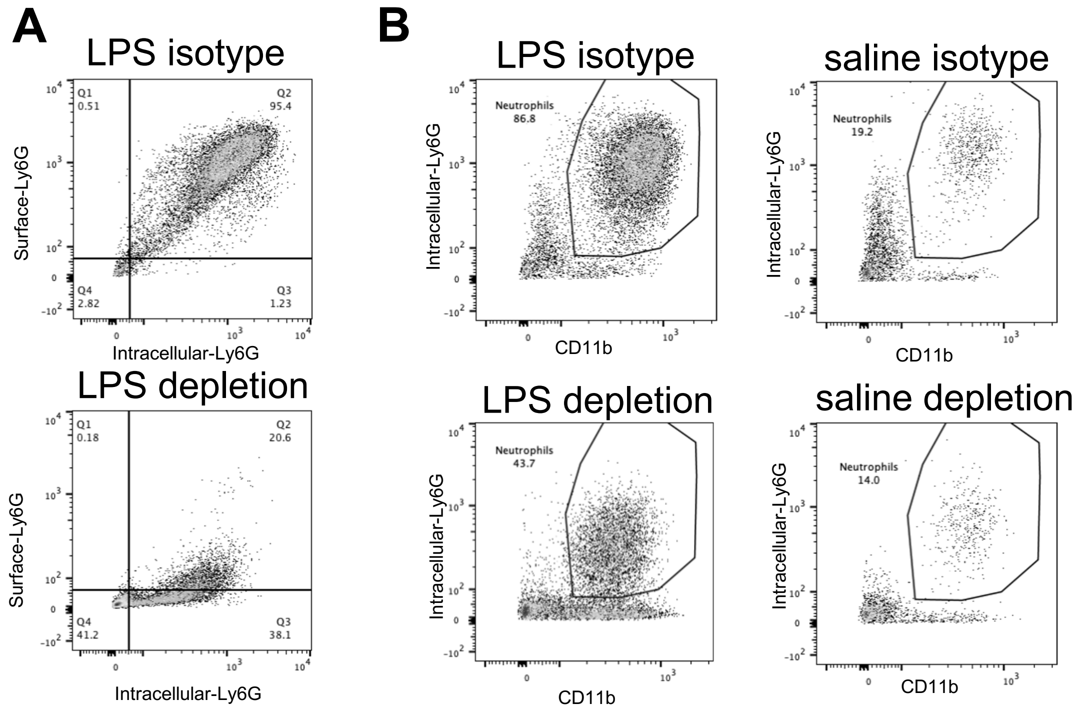

#### Supplementary Table 1: Purities of Isolated Cell Populations

Adherent Cell Populations (79.9 ± 0.14% $CD45^+$ cells)

| Sample | AMs | Mon./mac. | Neutrophils | Eosinophils | Lymphocytes ($CD45^+SSC\text{-}L^{lo}$) |
|---|---|---|---|---|---|
| **LPS** | 56.40% | 9.63% | 1.66% | 2.47% | 29.84% |
| **saline** | 69.40% | 3.23% | 0.24% | 0.50% | 26.63% |

Monocyte/macrophages for *in vitro* infections (> 99% $CD45^+$ cells)

| Sample | Mon./mac. | AMs | Neutrophils | Eosinophils |
|---|---|---|---|---|
| **LPS** | 95.9% | 0.26% | 2.88% | 0.72% |
| **saline** | 93.5% | 1.82% | 0.5% | 1.14% |

Neutrophils for *in vitro* infections (> 99% $CD45^+$ cells)

| Sample | Neutrophils |
|---|---|
| **LPS** | 98.7% |
| **saline** | 97.7% |

Percentages of total $CD45^+$ cells shown, assessed via flow cytometry. Data are each representative represent of 1 independent experiment with pools of 5 (adherent cells and monocyte/macrophages) or 2 (neutrophils) mice in each group.
